# Alzheimer’s Disease Neuropathological Comorbidities Are Common in the Younger-Old

**DOI:** 10.1101/2020.01.03.894451

**Authors:** Thomas G. Beach, Michael Malek-Ahmadi

## Abstract

Clinicopathological studies have demonstrated that Alzheimer’s disease dementia (ADD) is often accompanied by clinically undetectable comorbid neurodegenerative and cerebrovascular disease that alter the presence and rate of cognitive decline in aging and ADD. Aside from causing increased variability in clinical response, it is possible that the major ADD comorbidities may not respond to ADD-specific molecular therapeutics. As most reports have focused on comorbidity in the oldest-old, its extent in younger age groups that are more likely to be involved in clinical trials is largely unknown. We conducted a survey of neuropathological comorbidities in sporadic ADD using data from the US National Alzheimer’s Coordinating Center. Subject data was restricted to those with dementia and meeting National Institute on Aging-Alzheimer’s Association (NIA-AA) intermediate or high AD Neuropathological Change (ADNC) levels, excluding those with known autosomal dominant AD-related mutations. Subjects were divided into age-at-death categories for analysis: under 60, 60-69, 70-79, 80-89, 90-99 and 100 or over. Confirmatory of earlier reports, ADD histopathology is less severe with advancing age, effectively increasing the relative contribution of comorbidities, most of which rise in prevalence with age. Highly prevalent ADD comorbidities are not restricted to the oldest-old but are common even in early-onset ADD. The percentage of cases with ADD as the sole major neuropathological diagnosis is highest in the under-60 group, where “pure” ADD cases are still in the minority at 44%. After this AD as a sole major pathology in ADD declines to roughly 20% in the 70s and beyond. Comorbidity rates for some pathologies, especially LBD, are high even in subjects in their 60s and 70s, at nearly 60%, but for most others, their prevalence increases with age. TDP-43 pathology affects more than 35% of ADD subjects 80 and over while microscopic infarcts reach this rate a decade later. Gross infarcts rise more slowly and affect fewer subjects but still involve 15-20% of ADD after age 80. White matter rarefaction may be underestimated in the NACC database but is present in almost 70% of centenarians with ADD. Effective clinical trials depend on accurate estimates of required subject numbers, which are dependent on observed effect size and clinical response variability. Comorbidities are likely to affect both, leading to lower probability of clinical trial success. Stratifying ADD clinical trial analyses by presence and types of accompanying comorbidities might identify subgroups with higher effect sizes and greater clinical response rates, but accurate in-vivo diagnostic methods for most comorbidities are still lacking.

## Introduction

Neuropathological studies are increasingly demonstrating that Alzheimer’s disease dementia (ADD) is most often not the sole brain pathology in those that die and are autopsied, but is very likely to be accompanied by comorbid neurodegenerative and cerebrovascular disease. Comorbidities are likely to alter the presence and rate of cognitive decline in aging and ADD [1–40]. If all neuropathological changes found in elderly brains were considered, there would be few if any cases of “pure” ADD. While not all of these aging changes are likely be clinically significant [37], several of the commonly-observed ADD comorbidities have been demonstrated to contribute to cognitive impairment, including Lewy body disease (LBD) [4,23,26,30,41–47], hippocampal sclerosis [29,47–52], TDP-43 proteinopathy [12,13,37,53–57] and cerebral infarcts [1,10,11,30,35,46,57,58,58–70]. Aside from causing increased variability in clinical response, it is possible that these major comorbidities, each of which is individually capable of causing dementia, may not respond to ADD-specific molecular therapeutics because their molecular pathogenesis is different from ADD. At present, there are no accurate methods available with which to clinically diagnose these complications of ADD.

As the reported studies of comorbidities have focused on the oldest-old, and an increase in their prevalences with increasing age has been noted [19], it has been assumed by some that these would largely not be present in younger aged ADD subjects that might be more likely to be involved in clinical trials. However, in the Arizona Study of Aging and Neurodegenerative Disorders/Banner Sun Health Research Institute Brain and Body Donation Program [71], although the number of comorbid conditions increased with each decade, even subjects in their 70s had, on average, 2.8 major neuropathological diagnoses at autopsy [27]. To determine whether these findings were more broadly confirmed, we conducted a survey of neuropathological comorbidities in sporadic ADD using data from the National Alzheimer’s Coordinating Center (NACC)[72,73].

## Methods

We used a recent (10/09/19) NACC data download of all clinical and autopsy data with entries on NACC NP form version 10 [73]. The time period covered therefore is from January 2014 through August 2019. Included subject data was restricted to those with dementia and meeting National Institute on Aging-Alzheimer’s Association (NIA-AA) intermediate or high AD Neuropathological Change (ADNC) levels [74,75], considered to be a sufficient cause of cognitive impairment or dementia. Subjects known to have autosomal dominant AD-related mutations were excluded. Subjects were divided into age-at-death categories for analysis: under 60, 60-69, 70-79, 80-89, 90-99 and 100 or over.

Pathology categories investigated consisted of the major AD-specific lesions including senile plaques of all types, classified by Thal amyloid phase [76] (NACC variable NPTHAL), neurofibrillary tangles, classified by Braak stage stage [77,78] (NACC variable NPBRAAK), neuritic plaques, classified according to CERAD [79] (NACC variable NPNEUR), diffuse plaques, classified analogously to CERAD neuritic plaques (NACC variable NPDIFF), amyloid angiopathy, classified as none, mild, moderate or severe (NPAMY), and NIA-AA AD Neuropathological Change (ADNC) Level, with categories of intermediate or high [74,75](NACC variable NPADNC). Means and medians for each age category were compared. Decadal groups were compared using Kruskal-Wallis analysis of variance for ordinal AD pathology measures and chi-square tests were used to compare comorbidity prevalence rates.

Comorbid neurodegenerative conditions investigated included Lewy body disease (NACC variable NPLBOD; for this study only presence or absence in any brain region was recorded), hippocampal sclerosis (NACC variable NPHIPSCL; for this study recorded as present or absent); TDP-43 pathology (NACC variable NPTDPB; TDP-43 pathology present in amygdala), non-AD tauopathy (NACC variable NPFTDTAU; any non-AD tauopathy including progressive supranuclear palsy, corticobasal degeneration, Pick’s disease, argyrophilic grains, chronic traumatic encephalopathy, “tangle-only disease” or “other”). For each comorbid pathology and age category, the proportion of cases possessing that pathology was determined, relative to the number of cases for which the specific pathology type was assessed by contributing neuropathologists.

Comorbid cerebrovascular conditions investigated included circle of Willis arteriosclerosis (NACC variable NACCAVAS; for this study only “moderate” and “severe” qualified for presence of the condition), old gross cerebral infarcts (NACC variable NPINF; for this study 1 or more large or lacunar infarcts qualified for the condition), old microscopic infarcts (NACC variable NPOLD; for this study 1 or more old microinfarcts qualified for the condition), old microscopic hemorrhages (NACC variable NPOLDD; for this study 1 or more old microhemorrhages qualified for the condition), arteriolosclerosis (NACC variable NACCARTE; for this study only “moderate” and “severe” qualified for presence of the condition) and white matter rarefaction (NACC variable NPWMR; for this study only “moderate” and “severe” qualified for presence of the condition). For each comorbid pathology and age category, the proportion of cases possessing that pathology, relative to those for whom the pathology’s presence or absence was specifically recorded, was determined.

For all conditions, cases with missing data, or for which the specific condition was not assessed, were excluded.

## Results and Discussion

There were 1,839 cases that had dementia and met NIA-AA intermediate or high ADNC levels (Table 1). Confirmatory of earlier reports [19,52,80,81], ADD histopathology consistently becomes less severe with age (Table 1). It is important to realize that the decreased severity of AD histopathological lesions with age would effectively increase the relative importance of comorbidities in older subjects, most of which rise in prevalence with age.

**Table 1.**
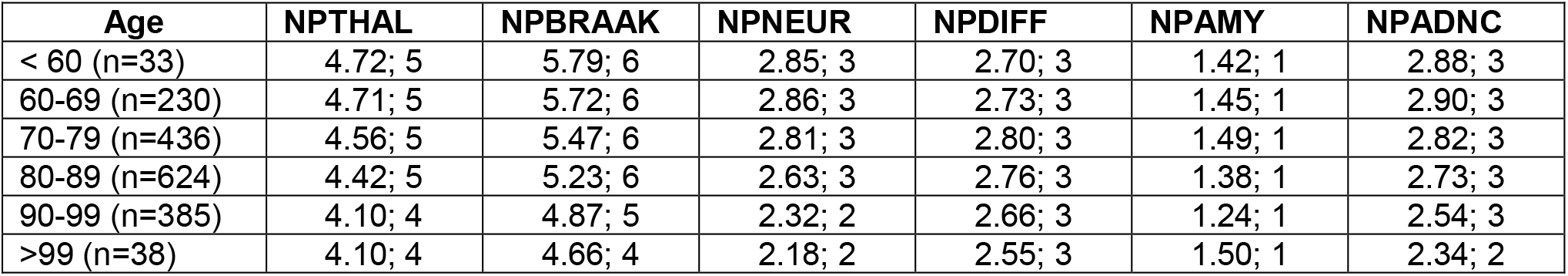
NACC autopsy data, by decade, for basic AD-related neuropathological variables in 1,839 dementia cases with intermediate or high ADNC. Number of cases, means and medians are shown. NPTHAL = Thal amyloid phase; NPBRAAK = Braak neurofibrillary stage; NPNEUR = CERAD neuritic plaque density; NPDIFF = diffuse plaque density; NPAMY = cerebral amyloid angiopathy density; NPADNC = NIA-AA AD Neuropathological Change level. Decadal groups are significantly different (p < 0.000001 for all AD lesion types except NPDIFF (p = 0.00046) and NPAMY (p = 0.0126).

Our analysis of these data shows that highly prevalent ADD comorbidities are not restricted to the oldest-old but are common even in early-onset ADD. Comorbidity rates for some pathologies, especially LBD, are high even in subjects in their 60s and 70s (Tables 2 and 3, Figure 1). The percentage of cases with ADD as the sole major neuropathological diagnosis, defined as the absence of LBD, TDP-43 pathology, non-AD tauopathy, hippocampal sclerosis, infarcts and microhemorrrhages, is highest in the under-60 group, where “pure” ADD cases are still in the minority, at 44%. After this AD as a sole major pathology in ADD declines to roughly 20% in the 70s and beyond.

**Figure 1.**
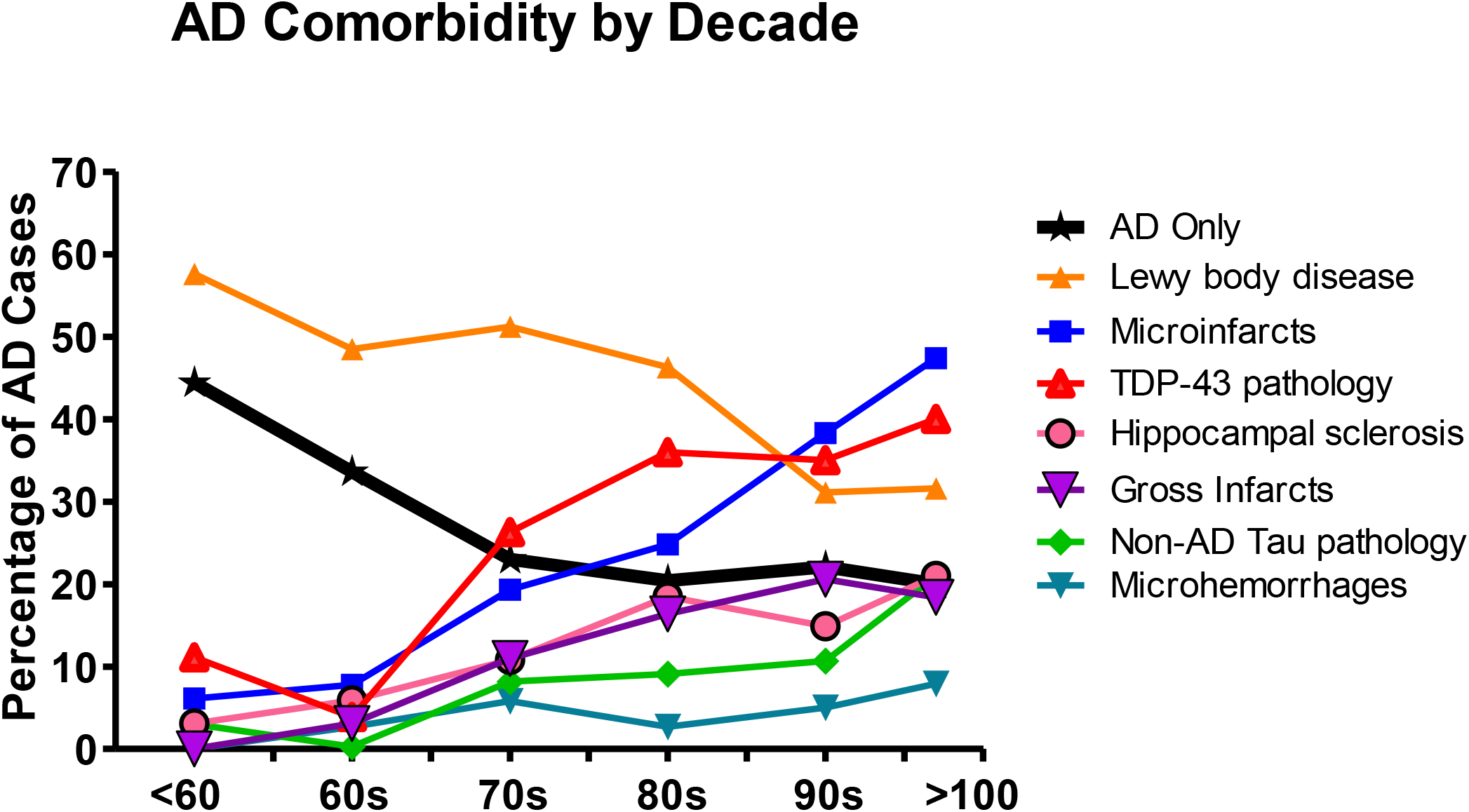
Major neuropathological comorbidities with ADD as compared to ADD as a sole diagnosis, by decade. The prevalences of all comorbidities shown increase with age, except for Lewy body disease.

**Table 2.** NACC data, by decade, for proportions and percentages of common neurodegenerative AD comorbidities in dementia cases with intermediate or high NIA-AA ADNC. Numerator is the number of cases meeting criteria for the comorbidity while denominator is the number of AD cases evaluated for the condition. AD Only = AD without any of the other diagnoses in this table and without NPINF, NPOLD and NPOLDD from Table 3; NPLBOD = Lewy body disease; NPHIPSCL = hippocampal sclerosis; TDPNOS = TDP-43 pathology at minimum present in amygdala; NPFTDTAU = non-AD tau pathology (PSP, CBD, Pick’s, argyrophilic grains, other). Decadal groups are significantly different (p < 0.001) except for AD Only (p = 0.02).

| Age | AD Only (n=835) | NPLBOD (n=1,737) | NPHIPSCL (n=1,800) | TDPNOS (n=1,006) | NPFTDTAU (n=1,812) |
| --- | --- | --- | --- | --- | --- |
| < 60 | 8/18; 44.4% | 19/33; 57.6% | 1/32; 3.1% | 2/18; 11.1% | 1/33; 3.0% |
| 60-69 | 40/119; 33.6% | 111/229; 48.5% | 19/320; 5.9% | 9/229; 3.9% | 10/320; 0.3% |
| 70-79 | 52/226; 23.0% | 221/432; 51.2% | 46/424; 10.8% | 66/251; 26.3% | 35/429; 8.2% |
| 80-89 | 63/308; 20.4% | 288/622; 46.3% | 113/610; 18.5% | 122/339; 36.0% | 56/617; 9.1% |
| 90-99 | 33/149; 22.1% | 119/383; 31.1% | 56/376; 14.9% | 54/154; 35.1% | 40/375; 10.7% |
| >99 | 3/15; 20% | 12/38; 31.6% | 8/38; 21.0% | 6/15; 40% | 8/38; 21.0% |

**Table 3.** NACC data, by decade, for proportions and percentages of common AD cerebrovascular comorbidities in dementia cases with intermediate or high NIA-AA ADNC. Numerator is the number of cases meeting criteria for the condition, denominator is the number of cases evaluated for the condition. NACCAVAS = severity of atherosclerosis of circle of Willis > mild; NPINF = gross infarcts including lacunes; NPOLD = old microinfarcts; NPOLDD = old cerebral microhemorrhages; NACCARTE = arteriolosclerosis > mild; NPWMR = white matter rarefaction > mild. Decadal groups are significantly different (p < 0.001) except for NPOLDD (p = 0.05).

| Age | NACCAVAS (n=1,707) | NPINF (n=1,832) | NPOLD (n=1,834) | NPOLDD (n=1,765) | NACCARTE (n=1,715) | NPWMR (n=1,657) |
| --- | --- | --- | --- | --- | --- | --- |
| < 60 | 1/33; 3.0% | 0/33; 0% | 2/33; 6.1% | 0/29; 0% | 6/32; 18.7% | 5/28; 17.9% |
| 60-69 | 39/319; 12.2% | 10/321; 3.1% | 25/322; 7.8% | 9/307; 2.7% | 77/309; 24.9% | 57/296; 19.3% |
| 70-79 | 145/317; 45.7% | 48/435; 11.0% | 84/435; 19.3% | 24/411; 5.8% | 176/413; 42.6% | 111/379; 33.7% |
| 80-89 | 275/616; 44.6% | 102/621; 16.4% | 154/622; 24.8% | 16/600; 2.7% | 326/577; 56.5% | 166/556; 29.9% |
| 90-99 | 225/384; 58.6% | 79/384; 20.6% | 147/384; 38.3% | 19/380; 5.0% | 209/352; 59.4% | 104/360; 28.9% |
| >99 | 27/38; 71.0% | 7/38; 18.4% | 18/38; 47.4% | 3/38; 7.9% | 19/32; 59.4% | 25/38; 65.8% |

Lewy body disease is the most common ADD comorbidity in subjects under 80, and in fact is most prevalent in AD subjects under 60, with its proportional presence gradually declining in older decades (Figure 1). As this closely parallels the decline in AD pathology severity with age, one conclusion is that LBD is largely synchronous or even dependent on AD pathology. The high reported rate of Lewy body disease in autosomal dominant ADD [82–89], and even in other cerebral amyloidoses [90,91] supports this conclusion. Prior work has consistently found that one-half or more of all subjects meeting clinicopathological diagnostic criteria for ADD also have LBD [71,92–94]. Similarly, up to one-half of subjects with dementia and Parkinson’s disease [95–107] and three-quarters or more of those with DLB, have clinically significant AD pathology [108–111]. In the great majority of subjects with both ADD and LBD, this co-existence is recognized only at autopsy [41,112–115], currently preventing, except for the minority with clinically-typical DLB, the exclusion or stratification of LBD subjects within ADD clinical trials.

The presence of LBD in ADD is clearly of clinical significance. Multiple autopsy-validated studies have indicated that cognitive decline is faster in ADD with any degree of associated LBD [26,42–45]. Formerly, it was not apparent whether or not this was primarily driven by dementia with Lewy bodies (DLB) subjects with neuropathologically-severe LBD (mostly neocortical stage diffuse Lewy body disease), as disease duration is reportedly shorter in this group [109,116]. The great majority of LBD in ADD subjects, however, does not meet neuropathological diagnostic criteria for DLB, due to insufficient pathology density and brain regional distribution [117–119]. These “AD-LB” cases are most often clinically silent [24] and diagnosed as probable ADD. We have recently reported that even in these AD-LB subjects without neocortical disease, as compared to ADD subjects without any LBD, that depression and Trail-Making Test A scores correlate significantly with LBD pathology, and the global rate of cognitive decline is more rapid than in either AD-DLB or ADD without LBD [41,120], even after adjustment for AD pathology severity. These data suggest that accompanying LBD is a “state” biomarker of a more clinically severe form of AD.

All the other ADD comorbidities increase in prevalence with age, suggesting that they are primarily age-dependent, or at least co-dependent on both AD and age (Figure 1). Of these, hippocampal sclerosis and TDP-43 pathology, which are thought to be closely related or even subsets of a single pathogenic class [50,80] are the next most common ADD comorbidity, at least up to about age 80-90, when TDP-43 pathology is present in about 35% of ADD cases (Table 2, Figure 1). Comorbid TDP-43 has been reported by several groups to contribute to cognitive impairment in ADD [12,13,37,53–57].

Cerebrovascular disease, as a whole, may be of equal or greater importance as an ADD comorbidity, as compared to LBD and TDP-43. Microscopic infarcts rise steadily in prevalence with age, at 40-50% becoming the most common ADD comorbidity after age 90. Gross infarcts rise more slowly and affect fewer subjects but still involve 15-20% of ADD after age 80. Both of these infarct classes have been reported to be independent influences on cognition [1,10,11,30,35,57–69]. White matter rarefaction (WMR) has long been reported to be roughly twice as common in ADD as compared to normal elderly, being present in 60% or more of ADD subjects [121–123]. At 30-70%, the NACC database may be underestimating WMR. This may be due to the difficulty of detecting it with only standard small-block histology sampling. Without setting thresholds of severity, MRI-detected WMR has been reported to be present in 39-96% of elderly subjects and close to 100% of those with ADD [124]. Correlations of WMR with cognition have been complicated by co-linearity with AD histopathology as well as a dual vascular and AD-related pathogenesis [58,70,125–132], but when WMR rarefaction in ADD is severe, or in non-demented older subjects where AD histopathology is minimal, there are significant associations, especially with frontal lobe functions or longitudinal decline [124,133–136]. Like LBD, WMR is highly prevalent in autosomal dominant early-onset AD and thus is probably at least partially dependent on AD-related factors [137–139].

In the NACC database, microhemorrhages are much less common than infarcts and WMR, reaching only 5-8% prevalence after age 80. Again these may be underestimated at autopsy as compared to MRI, or when special efforts are made in postmortem examination, as one autopsy center has reported a prevalence rate of 62% in oldest-old subjects [140]. In the Alzheimer’s Disease Neuroimaging Initiative, 25% of a mixed group of subjects, with normal cognition, mild cognitive impairment, and ADD, had microhemorrhages, with a tendency for proportionately more in AD [141]. Microhemorrhages are reported to increase risk for incident dementia [142].

Non-AD tauopathies, with microscopic tau pathologist that is morphologically unlike the neurofibrillary tangles of AD, are relatively uncommon ADD comorbidities in the NACC database, being present in under 10% or less of subjects under 90. It is possible that these tauopathies may often be overlooked against the background of severe AD neurofibrillary pathology. Reports from centers that specifically look for argyrophilic grains and aging-related tau astrogliopathy (ARTAG) find AD comorbidity rates up to 40% for both [143–150]. The classical conditions in this group, including progressive supranuclear palsy (PSP), corticobasal degeneration (CBD) and Pick’s disease, are much less prevalent, although PSP may be considerably more common than previously realized due to low clinical sensitivity for the diagnosis [143,151–158].

Effective clinical trials depend on accurate estimates of required subject numbers and variability as well as effect size of the treatment. Increased variability of clinical decline rates will require larger subject numbers to overcome. Effect size depends on the efficacy of the therapeutic agent, which may be at least in part dependent on a matching of molecular mechanisms of agent and pathogenesis. Neuropathological comorbidities in AD affect rates of cognitive decline and are by definition different in their molecular pathogenesis from AD. Comorbidities are thus likely to lead to lower probability of clinical trial success. Stratifying AD subjects by presence and types of accompanying comorbidities might result in increased observed effect sizes in some groups as compared to others, potentially “rescuing” failed clinical trials. Accurate in-vivo diagnostic methods are urgently needed for the major AD comorbidities.

## Acknowledgements

The NACC database is funded by NIA/NIH Grant U01 AG016976. NACC data are contributed by the NIA-funded ADCs: P30 AG019610 (PI Eric Reiman, MD), P30 AG013846 (PI Neil Kowall, MD), P30 AG062428-01 (PI James Leverenz, MD) P50 AG008702 (PI Scott Small, MD), P50 AG025688 (PI Allan Levey, MD, PhD), P50 AG047266 (PI Todd Golde, MD, PhD), P30 AG010133 (PI Andrew Saykin, PsyD), P50 AG005146 (PI Marilyn Albert, PhD), P30 AG062421-01 (PI Bradley Hyman, MD, PhD), P30 AG062422-01 (PI Ronald Petersen, MD, PhD), P50 AG005138 (PI Mary Sano, PhD), P30 AG008051 (PI Thomas Wisniewski, MD), P30 AG013854 (PI Robert Vassar, PhD), P30 AG008017 (PI Jeffrey Kaye, MD), P30 AG010161 (PI David Bennett, MD), P50 AG047366 (PI Victor Henderson, MD, MS), P30 AG010129 (PI Charles DeCarli, MD), P50 AG016573 (PI Frank LaFerla, PhD), P30 AG062429-01(PI James Brewer, MD, PhD), P50 AG023501 (PI Bruce Miller, MD), P30 AG035982 (PI Russell Swerdlow, MD), P30 AG028383 (PI Linda Van Eldik, PhD), P30 AG053760 (PI Henry Paulson, MD, PhD), P30 AG010124 (PI John Trojanowski, MD, PhD), P50 AG005133 (PI Oscar Lopez, MD), P50 AG005142 (PI Helena Chui, MD), P30 AG012300 (PI Roger Rosenberg, MD), P30 AG049638 (PI Suzanne Craft, PhD), P50 AG005136 (PI Thomas Grabowski, MD), P30 AG062715-01 (PI Sanjay Asthana, MD, FRCP), P50 AG005681 (PI John Morris, MD), P50 AG047270 (PI Stephen Strittmatter, MD, PhD).

